# BREVIPEDICELLUS and ERECTA mediate expression of *AtPRX17* in preventing Arabidopsis callus retardation and browning

**DOI:** 10.1101/2021.02.18.431912

**Authors:** Junyan Xie, Bin Qi, Yuanyuan Wu, Chenghong Mou, Lihua Wang, Yuwei Jiao, Yanhui Dou, Huiqiong Zheng

## Abstract

Efficient in vitro callus generation is fundamental to tissue culture propagation, a process required for plant regeneration and transgenic breeding for desired phenotypes. Identifying genes and regulatory elements that prevent callus retardation and browning is essential to facilitate the development of vitro callus systems. Here we show that *BREVIPEDICELLUS* (*BP*) and *ERECTA* (*ER*) pathways in *Arabidopsis* callus are converged to prevent callus browning and positively regulate an isoperoxidase gene At*PRX17* expression in the rapid growth callus. Loss of functions in both *BP* and *ER* resulted in markedly increasing callus browning. Transgenic lines with *pro35S*::*AtPRX17* in the *bp-5 er105* double mutant background fully rescued this phenotypic abnormality. Using plant *in vitro* DNA-binding assays, we observed that BP protein bound directly to the upstream sequence of *AtPRX17* to promote its transcription during callus growth. ER is a universally presenting factor required for cell proliferation and growth, we show that *ER* positively regulates expression of a transcription factor *WRKY6*, which also directly binds to an additional site of the At*PRX17* promoter for its high expression. Our data reveals an important molecular mechanism in regulating expression of peroxidase isozyme to reduce Arabidopsis callus browning.

**Highlight:** *BREVIPEDICELLUS* and *ERECTA* are involved in regulating Arabidopsis callus browning by controlling expression of *AtPRX17*.

## Introduction

Callus retardation and browning are major impediments in vitro culture of plants, resulting in decreased regenerative ability, poor growth, and even death. Oxidative stress has been considered as one of major factors in induction of callus retardation and browning. During plant tissue culture, abundant ROS could accumulate in the rapid proliferation cells because of the high metabolic rate (Cosio and Dunand, 2009; Wells et al., 2010). The accumulation of ROS not only significantly slows the further cell proliferation rate, but often causes the callus browning (Laukkanen et al.,1999). However, in the natural conditions, tissues of higher plants, such as shoot and root meristems, usually experience a long-time continued and relatively fast cell division, whereas they are barely disturbed by cell browning even if the metabolic process always produces more ROS (Hirt and Apel, 2004; Wells et al., 2010; Tsukagoshi et al., 2010; Ikeuchi et al., 2013). One possibility is that plants could possess an efficient mechanism to scavenge ROS to ensure cell proliferation at a decided rate, however, very little is known about the molecular mechanism(s) for plant to eliminate the excess ROS in both meristematic tissues *in vivo* and developing callus tissues *in vitro*. The KNOTTED1-like HOMEOBOX (KNOX) transcription factor BREVIPEDICELLUS (KNAT1/BP) is involved in maintaining meristem of shoot apexes (Smith et al., 2002), xylem fiber development (Woerlen et al., 2017; Felipo-Benavent et al., 2018; Milhinhos et al., 2019) and fruitlet abscission (Zhao et al., 2020). Overexpression of the maize KNOTTED1 gene caused a switch from determinate to indeterminate leaf cell fates (Sinha et al., 1993). In the process of meristematic growth, the continued and rapid cell proliferation occurs in some special zones, such as the peripheral zone in shoot apical meristem and the meristem zone adjacent to the root stem cells, in which *BP* plays a role in part by spatially regulating boundary genes (Woerlen et al., 2017). Down-regulated expression of a peroxidase encoded by At*PRX9GF* in *bp-9* seedlings was reported (Mele et al., 2003). In addition, BP is involved in regulating expression of *SOBIR1/EVR* gene, which acts together with ERECTA (Terpstra et al., 2010) to regulate programmed cell death (PCD) during xylem development in Arabidopsis shoot (Milhinhos et al., 2019). Reactive oxygen species (ROS) are emerging as intracellular signaling molecules that efficiently regulate PCD during xylem development and cell proliferation in meristem in shoots and roots (Breusegen and Dat, 2006; Tsukagoshi et al., 2009). Whether BP is involved in regulation of ROS signaling remains unknown.

Peroxidase catalyzes the reduction of H_2_O_2_ and protects tissues and cells from oxidative damage and plays important roles in controlling callus growth and browning (Brown *et al*., 1993; Creissen *et al*., 1994; Wakui *et al*., 1999; Zhang *et al*., 2020). However, apparently conflicting roles of peroxidase in the initiation of meristematic activity as well as with the suppression of growth on callus development were reported in previous studies (Basile, 1980; Goff 1975; Habib et al., 2014). An explanation on those inconsistent results is attributed to the occurrence of peroxidase in multimolecular forms, *i*.*e*. isoenzymes. Individual isoperoxidases may differ in their substrate specificity, pH optima, and distribution within cellular compartments, etc. (Kay et al., 1987). Specific isoperoxidases have been proven to be correlated with specific developmental events (Cosio and Dunand., 2009). Nevertheless, no specific isoperoxidase has been identified, isolated, and characterized with respect to its possible role in controlling callus growth and browning.

In this work, we report that a preoxidase gene At*PRX17* plays a critical role involved in preventing callus browning. At*PRX17* belongs to the class III peroxidase family (EC 1.11.1.7) (Tognolli et al., 2002), and we show that upregulation of At*PRX17* is directly promoted by two types of transcription factors: one is the meristem-specific regulatory protein BP (Serikawa et al., 1996; Hay and Tsiantis, 2010), and the other is WRKY6 (Eulgem and Somssich, 2007), a transcription factor that is positively regulated by the putative receptor protein kinase ERECTA (ER).

## Materials and methods

### Plant materials and culture conditions

*Arabidopsis thaliana* wild-type ecotype Columbia (Col), Landsberg *erecta* (L*er*) and Landsberg *ERECTA* (Lan), mutant *brevipedicellus* (*bp*) *bp-1* and *bp-5* (Douglas et al., 2002; Qi and Zheng, 2013), *er-105* (Hord et al., 2008) and transformants *proBP::GUS* (Ori et al., 2000) and pro 35S::BP (Lincoln et al., 1994) have been previously described. *prx17* (SALK_034684c) and *wrky6* (SALK_012997c) are (T)-DNA insertion alleles in the Columbia (Col) accession and was obtained from the Arabidopsis Biological Resource Center. 3∼5 mm root segments from 7-d old *Arabidopsis thaliana* seedlings were cut and transferred to callus induction medium (CIM): MS medium (Murashige and Skoog, 1962) with 3% sucrose, 2 mg/L 2,4-dichlorophenoxyacetic acid, and 0.8% agar. Explants were incubated on CIM for 5 weeks at 25°C under dark conditions, then calli produced from the explants was subcultured on new CIM for twice. The suspension cultures for these calli were grown as described (Zhang et al., 2015). The suspension cultures were taken as stock for repeated callus formation. The cultures were then subcultured on solid medium again and callus with a diameter of about 1mm (about 7d after subculture on the new medium) was used to monitored the growth rate by weighed the fresh weight of cells.

### Scanning electron microscopy

Samples were prepared as described by Wei et al. (2010). Briefly, calli that had been subcultured for 21 days was fixed in 3% glutaraldehyde-phosphate buffer saline fixative solution (pH 7.2) for overnight, then the samples were mounted on aluminum stubs and coated with gold in JEOL JFC-1600 after graded dehydration and replacement. The treated calli was observed with JEOL JSM-6360LV scanning electron microscope at 6kV.

### Transmission electron microscopy

Samples were prepared as described by Bestwick et al. (1997). Briefly, calli as described above was incubated in 5 mM CeCl_3_ in 50 mM 3-(N-morpholino) propanesulfonic acid (Mops) at pH7.2 for 1h. After treatment, samples were fixed in 1.25%(v/v) glutaraldehyde and 1.25%(v/v) paraformaldehyde in 50 mM sodium cacodylate (CAB) buffer, pH7.2 for 1h and post-fixed in 1%(v/v) osmium tetroxide in CAB for 45 min. After being dehydrated through an ethanol series, samples were infiltrated and embedded in Epon 812 resin. The specimens were sectioned and the thick section (0.5μm) were stained with 0.1% toluidine blue and examined under light microscopy (Leica DMLB). The ultra-thin sections (about 100 nm) were examined using a transmission electron microscope (Hitachi 7650 TEM) at an accelerating voltage of 75kV.

### Diaminobenzidine oxide and nitroblue tetrazolium staining

In situ detection of hydrogen peroxide was performed by staining with diaminobenzidine oxide (DAB) using an adaptation of a previous method (Daudi et al., 2012). Briefly, calli that had subcultured for 21 days was stained for 2h in 1mg/ml DAB solution containing Tween 20 (0.05% v/v) and 10 mM sodium phosphate buffer (pH7.0) in a Eppendorf tube. The staining was terminated in ethanol: glycerol: acetic acid 3:1:1(bleaching solution) placed in a water bath at 95°C for 15 min. For nitroblue tetrazolium (NBT) staining, calli was stained for 15 min in a solution of 2mM NBT in 20 mM phosphate buffer pH6.1. The reaction was stopped by transferring the calli in distilled water. At least six independent culture plates were used as biological replicates and three callus blocks were sampled from each culture plate.

### TUNEL assay

The calli was fixed with 4%(w/v) fresh paraformaldehyde in PBS and labeled by the TUNEL reaction mixture: the terminal deoxynuleotidyl transferase solution and the label solution described in the kit’s manual. The stained cells were analyzed with a fluorescence microscope (Olympus, FV1000). The excitation and emission wave lengths were 488 nm and 515 nm, respectively.

### Evans blue staining

Cell death in calli was assayed as described by Baker and Mock (1994). Briefly, a 50 mg callus was added to a mixture of 0.5 ml of 1% (w/v) Sucrose and 0.5 ml of 0.5%(w/v) Evans blue solution. After 10 min, cells were drained and rinsed with deionized water, 30-40 ml, until no further blue eluted from the cells. After washing, the cells gently transferred to a 1ml plastic Eppendorf tube containing carborundum (<0.1mg). 0.5 ml of 1% aqueous SDS was added to cells to release the trapped Evans blue from the cells. The cells were ground and the homogenate diluted with 0.5 ml of deionized water, centrifuged at 10,000×g for 3 min. A 0.8 ml aliquot of the supernatant was removed and the optical density determined spectrophotometrically at 600 nm.

### Peroxidase isoenzyme analysis

The classical guaiacol peroxidase (class III peroxidases, E. C. 1.11.1.7) isoenzymes were analyzed as described by Naton et al. (1992). Briefly, three-week subcultured callus was homogenized on ice in 50 mM Tris-HCl buffer (pH 7.4) containing 0.58 mol/L sucrose. The extracts were centrifuged (10,000 g, 20 min, 4°C) and the supernatant collected as the peroxidase fraction. For enzyme visualization, the procedures of Brewbaker et al. (1968) were modified. Peroxidases were visualized by adding 4 ml substrate solution (2% 3, 3′-diaminobenzidine tetrahydrochloride in 0.1mM Tris-acetate buffer, pH 4.5) to 15.2 ml dd H_2_O, and starting the reaction with 0.8 ml 3%H_2_O_2_ (v/v). Incubation for 5∼15 min at room temperature revealed greenish-brown bands on a light-yellow background.

### Reverse transcription-polymerase chain reaction analyses (RT-PCR) and qRT-PCR

Total RNA was extracted from three-week subcultured callus using an RNAiso plus Kit (TaKaRa, http://www.takara.com.cn) according to the manufacturer’s instructions. cDNA was synthesized using 2 μg of total RNA and 100 U of ReverTra Ace reverse transcriptase (Toyobo Co., Ltd, Japan) according to the manufacturer’s instructions. The products were subsequently taken to amplify the targeted genes with primers for *PRX17*; *PRX52*; *WRKY6*; *WRKY15*; *WRKY 25*; *WRKY 33* and *WRKY46*. The constitutive housekeeping gene *ADENINE PHOSPHORIBOSYL TRANSFERASE1* (*APT1*) was used as an internal control. Reactions of qRT-PCR were done in a 384-well plate format with 7900HT Fast Real-time PCR System (Applied Biosystems®, http://events-na.appliedbiosystems.com), and SYBR Green to monitor double-stranded DNA synthesis. The *APT1* gene was used as reference for the *BP, ER*, At*PRX17* and *WRKY6* genes. Experiments were conducted at least three times with equivalent results. The primers used in this study are listed in Supplemental Table S1.

### Screening and isolation of the Arabidopsis T-DNA insertion mutants, *prx17* and *wrky6*

For identification of homozygous insertion of *prx17* and *wrky6* mutants, respectively, segregation analysis was performed by genotyping progeny of the mutant lines containing T-DNA insertion in theAt*PRX17*gene (SALK_034684c) or in *WRKY6* gene (SALK_012997) in various crosses. Progeny seedlings (about 4-week-old) were genotyped by extracting DNA from a single leaf of the seedling using PCR genotyping as described by Shpak et al. (2003). RT-PCR was performed to verify knock-out of theAt*PRX17*and *WRKY6* transcript. The primers used in this study are listed in Supplemental Table S1.

### *35S::PRX17* transgenic plants

A complementary vector consisted of CaMV35S promoter fused to the *PRX17*-encoding genomic fragment (Col-0, wild-type) was prepared according to Venglat et al. (2002). Briefly, a DNA fragment containing the full-length genomic coding region for At*PRX17* gene was amplified by PCR using the Col-0 cDNA template and the primers: 5’-<u>CTGCAG</u>ATGTCTCTTCTTCCCCAT-3’ and 5’-<u>GAGCTC</u>TCAAGATACAAGCAATAC-3’. The amplified fragments were digested with *PstI* and *SacI* and inserted into the *PstI* and *SacI* sites of pMD18-T-Vector. Subsequently, the *PstI/SacI*-amplified fragment was digested by *PstI/SacI* and ligated to CaMV 35S promoter of binary vector pHB. *Agrobacterium tumefaciens* (GV3101) containing this recombinant construct was used to transform *bp-5 er-105* plants as described by Clough and Bent (1998). All transgenic lines used in this study are T3 homozygous plants with single copy insertion.

### Electrophoretic mobility shift assay (EMSA)

For synthesis and purification of recombinant BP and WRKY6 proteins, cDNAs containing the full-length coding region of these two proteins were amplified by PCR. The amplified *BP* or *WRKY6* cDNA was cloned into the vector pET-30a (Novagen, www.novagen.com) by using *BamHI* and *Sacl* sites. The PCR primers for the *BP* and *WRKY6* amplification were listed in SupplementalTable S1.

All constructs were verified by sequencing. The 6xHis-BP and 6xHis-WRKY6 expression plasmid was transformed into the bacterial strain BL21 (DE3) pLysS. The transformed cells were cultured at 37°C until the OD_600_ of the cell culture was 0.5, and then induced with 1 mM IPTG for 36 h at 12°C. For extraction of native fusion protein, the cultured bacteria cells were lysed by using a high-pressure cell crusher and the fusion proteins were purified with Ni-NTA resin (Qiagen, www.qiagen.com) according to the manufacturer’s instructions.

For EMSA, the complementary pairs of biotin-labeled oligonucleotides corresponding to the At*PRX17* promoter region containing BP or WRKY6 binding sites was obtained by PCR amplification using 5’-biotin-labeled primers (Sangon Biotech, Shanghai Co., Ltd) to generate double-stranded probes. DNA binding reactions were performed with the Light Shift Chemiluminescent EMSA Kit (Pierce, www.piercenet.com) according to the instructions. DNA binding reactions were performed in a total volume of 20 μl of buffer (10mM Tris-HCl, pH 7.5, 2.5% Glycerol, 50mM KCl, 1mM DTT, 5mM MgCl_2_) containing 50ng/μL poly[dI-dC], 0.05% NP-40, 1 μg of the recombinant His-tagged BP or WRKY6 proteins and 20 fmol of probe. The binding specificity was assessed by competition with a 100- or 200-fold excess of unlabeled double-stranded oligonucleotides. Binding reaction mixtures were incubated for 30 min at 23°C and separated on native PAGE gels (5% polyacrylamide gel) in 0.5×TBE buffer, at 100V for 90 min. After electrophoresis, gels were bolted onto a positively charged nylon membrane (Amersham, now GE Healthcare, http://www.gelifesciences.com). The DNA was linked using a UV light cross-linker instrument equipped with 254-nm bulbs for 0.8 min exposure.

### Transient GUS assay by agroinfiltration of *Nicotiana benthamiana*

*PRX17* promoter sequence was amplified with specific primers (the forward primer 5’-AAGCTT TGGGACTGAATGAAACTGCTGA-3’ and the reverse primer 5’-GGATCCACTTTTTTCTTTTTTGGTGTTG-3’) by PCR from Arabidopsis genomic DNA and cloned into the transformation vector pCAMBIA1300-pBI101 at the HindIII and BamHI restriction sites, respectively as Reporter *PRX17*_*pro*_::GUS. The *BP* full-length cDNA sequence was amplified with *BP*-specific primers (the forward primer 5’-GAATTCATGGAAGAATACCAGCATGACAACAG-3’ and the reverse primer 5’-GTCGACTTATGGACCGAGACGATAAGGTCCAT-3’) cloned into pC1300-N1-YFP vector at EcoRI and SalI sites as Effector 35S-BP. The *WRKY6* full-length cDNA sequence was amplified with *WRKY6*-specific primers (the forward primer 5’-GGATCCATGGACAGAGGATGGTCTGGTCTCA-3’ and the reverse primer 5 ‘-GTCGACTTGATTTTTGTTGTTTCCTTCGC-3’) and cloned into pC1300-N1-YFP vector at BamHI and SalI sites as Effector 35S_*pro*_::WRKY6. The constructs of *PRX17*_*pro*_::GUS with 35S_*pro*_::BP or 35S_*pro*_::WRKY6, were transformed into Agrobacterium (GV3101). Agrobacteria was infiltrated into intact leaves of *Nicotiana benthamiana* as previously described (Kane et al., 2007). After infiltration, plants were kept at 23°C for 3 days. Histochemical GUS assay was performed as previously described (Wei et al., 2010).

### Chromatin Immunoprecipitation (ChIP)

ChIP assay was performed using a method modified from the Chromatin Immunoprecipitation (ChIP) Assay Kit (Upstate,Catalog # 17-295). Callus was incubated in 1% formaldehyde for 30 min under vacuum. The cross-linking was stopped by adding glycine to a final concentration of 0.125 M. Tissues were rinsed with water and ground into a fine powder with liquid nitrogen. To extract chromatin, the powder was resuspended in SDS lysis buffer (1% SDS, 10 mM EDTA, 50 mM Tris, pH 8.1, 1 mM phenylmethylsulfonyl fluoride (PMSF)). The chromatin DNA was sonicated to reduce DNA length and diluted 1:10 in chromatin immunoprecipitation dilution buffer (0.01% SDS, 1.1% Triton X-100, 1.2 mM EDTA, 16.7 mM Tris-HCl, pH 8.1, 167 mM NaCl). The chromatin solution was precleared with Protein A agarose beads blocked with salmon sperm DNA (Upstate Biotechnology). Immunoprecipitations were performed with anti-BP antibody (sc-19215). The BP-bound chromatin was purified by incubation with Protein A agarose beads blocked with salmon sperm DNA and washing with low-salt wash buffer (0.1% SDS, 1% Triton X-100, 2 mM EDTA, 20 mM Tris-HCl, pH 8.1, 150 mM NaCl), high-salt wash buffer (0.1% SDS, 1% Triton X-100, 2 mM EDTA, 20 mM Tris-HCl, pH 8.1, 500 mM NaCl), LiCl wash buffer (0.25 M LiCl, 1% IGEPAL-CA630, 1% deoxycholic acid (sodium salt), 1 mM EDTA, 10 mM Tris, pH 8.1), and TE buffer. The BP-bound chromatin was eluted from Protein A agarose beads with elution buffer (1% SDS, 0.1 M NaHCO_3_). After reversing the cross-linking by the addition of NaCl to a final concentration of 200 mM and incubation at 65°C for 4h, the DNA was purified by treating with 40 mg/mL proteinase K for 1 h, following by phenol/chloroform extraction and precipitation with DNA mate (TaKaRa). The primers used for PCR were listed in Supplemental Table S1.

## Results

### Combined action of *BP* and *ER* in controlling callus growth

In the wild-type (Col), root tips as explants from 7-day-old seedlings on callus induction medium (CIM) are usually able to generate the earliest callus at about 3th day after culture (DAC). The callus then undergoes a rapid growth from 10 DAC to 24 DAC (Fig. 1A and E, Fig. S1A-F), accompanied by rapidly increased *BP* expression in new regeneration callus (Fig. S1G-J). The first appearance of calli from the same-age root explants of *bp-5* and *er-105* mutants as well as *bp-5 er-105* double mutant (Col background) was also at about 4 DAC and the callus grew normally as wild-type (Fig. S2). However, callus of double mutant *bp5 er105* showed significantly retarded growth rate and browning appearance after 3-4 times subculture (Fig. 1D), while *bp-5* and *er-105* callus grew normally as their wild-type (Fig. 1A, B and C). Compared with those of Col, *bp-5*, and *er-105*, the total fresh weight produced by the *bp-5 er-105* explants was significantly reduced (Fig. 1E). In addition, we also analyzed another loss-of-function mutant of *BP* (*bp-1*), which is in the Landsberg *erecta* (L*er*) genetic background, and similar to the *bp-5 er-105* double mutant, callus growth of *bp-1* exhibited a markedly slowed manner in comparison with calli of *bp/ER* mutant, L*er* (*BP*/*er*) and Lan (*BP/ER*) (Fig. S3). These results indicated that *BP* and *ER* could co-regulate the growth and browning of Arabidopsis callus.

**Fig. 1.**
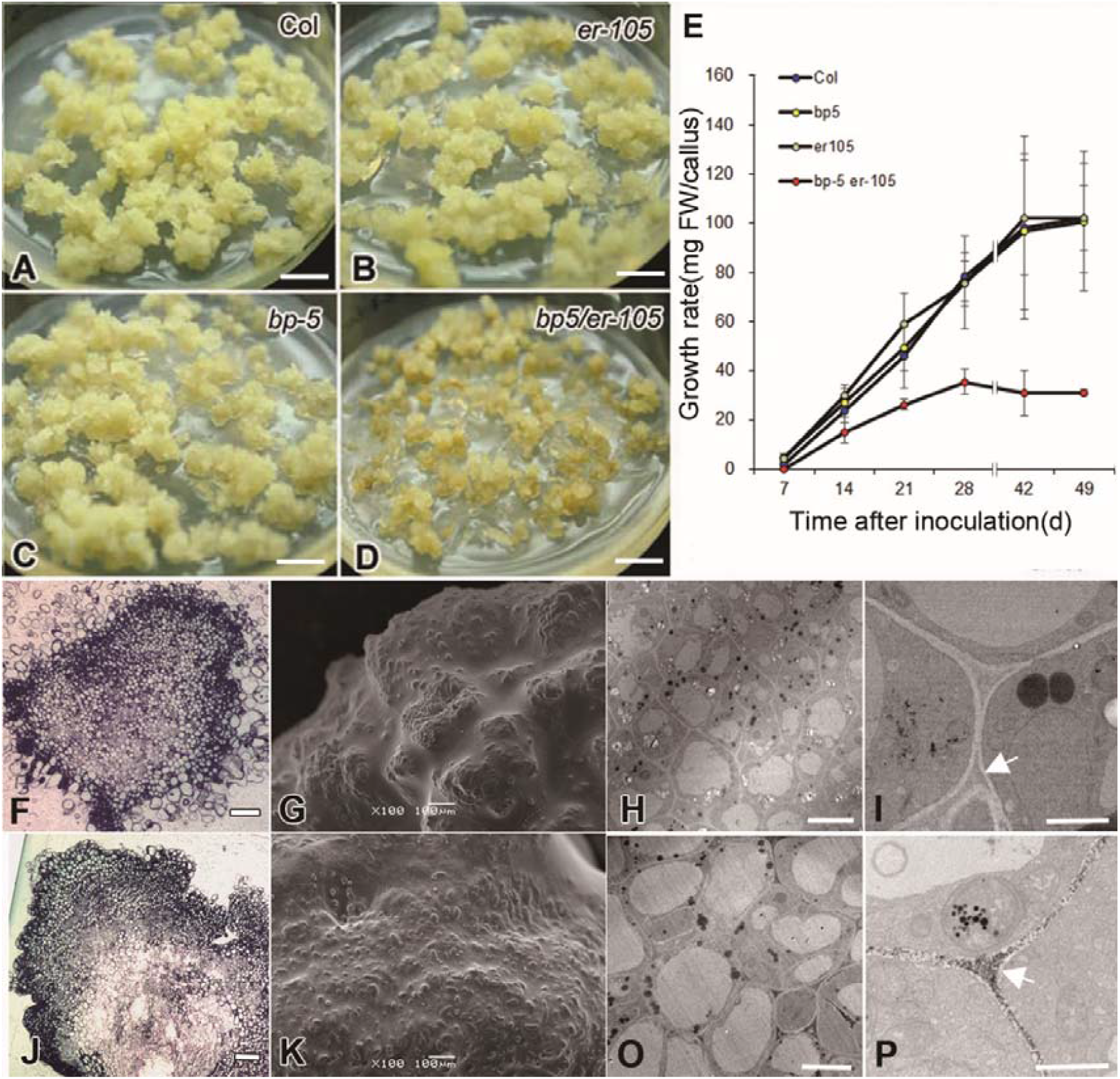
*BP* and *ER* genes implicate in callus growth and browning. **(A-D)** Callus of wild-type Col (A), single mutant er-105(B) and bp5 (C) and double mutant bp5 er-105 (D) after 3 weeks cultured on MS medium. Note that double mutant bp5 er-105 callus appears smaller and oxidative browning (D). Scale bars, 10 mm. **(E)** The growth rate of culture cells among the double mutant (bp-5 er-105), single mutant (er-105 and bp-5) and Col. Error bars indicate the Standard deviation of the mean for three replications(n≥20 calli for each time point). **(F and J)** Resin sections of Col (F) and bp5 er-105 (J) callus. Scale bars, 100μm. **(G and K)** Scanning electron microscopy images of Col (G) and bp5er-105 (K) callus. Scale bars, 100μm. **(H and O)** Electron micrographs of sections of Col (H) and *bp5 er-105* (O) callus, which was stained by 10 mM cerium chloride (CeCl_3_) to compare the localized H_2_O_2_ in cells between Col (H) and *bp5 er-105* (O) calli. Twenty calli from each genotype was collected for analysis of resin sections and electron microscopy. Scale bars, 20μm. (I) and (P) are enlarged from (H) and (O), respectively. Arrows point that the deposits are formed throughout cell wall of *bp5 er-105* callus cells, but less in cell wall of Col callus cells. Scale bars, 2μm.

### Hydrogen peroxide and superoxide in relation to BP/ER controlling callus growth

The histological examination indicated that the cell arrangement of *bp-5 er-105* calli was smaller and denser in the outer layer, but flaccid in the central region (Fig. 1J) compared to its wild-type Col (Fig. 1F). Observations using scanning electron microscopy (SEM) revealed that Col calli consisted of many globular nodules (Fig. 1G), whereas *bp-5 er-105* calli had fewer globular nodules (Fig. 1K). Several previous studies indicated the role of ROS homeostasis in the callus development (Tang et al., 2004; Che et al., 2006; Zhang et al., 2018). To test whether the distribution of H_2_O_2_ in *bp-5 er-105* callus cells was different from that in wild-type, we used the cerium chloride assay, in which cerium chloride forms electron-dense precipitates in the present of H_2_O_2_ through formation of cerium perhydoxide (Bestwick et al., 1997; Shen et al., 2015). With this assay, we observed apparently localized cerium precipitations in the *bp-5 er-105* callus cells (Fig 1O and P), especially in the cell wall (Fig 1P), in comparison with that of wild-type Col callus cells (Fig. 1H and I). This result indicated highly accumulation of H_2_O_2_ in cells of *bp-5 er-105* double mutant during rapid growth.

High level of H_2_O_2_ could induce programmed cell death (PCD), which can be judged based on DNA fragmentation using the terminal deoxynucleotidyl transferase dUTP nick end labeling (TUNEL) assay (Biswas and Mano, 2015). More than 50% of the cells of 21 DAC *bp-5 er-105* callus had positive TUNEL staining, while less than 20% of the cells of wild-type (Col) and two single mutants *bp-5* and *er-105* callus displayed positive TUNEL staining (Fig. 2A). The ratio of death cells in *bp-5 er-105* callus was apparently higher than those of Col, *bp-5* and *er-105* callus, as detected by Evans blue (Fig 2B). To understand whether the defective callus growth and cell death of *bp-5 er-105* is related to the accumulation of H_2_O_2_ in the cells, we analyzed H_2_O_2_ and superoxide (O_2_^.-^) levels in *bp-5 er-105* callus by comparing with those in Col, *bp-5* and *er-105* callus, respectively. H_2_O_2_ level in *bp-5 er-105* callus was apparently higher than those in Col, *bp-5* and *er-105* callus (Fig. 2D), whereas the O_2_. ^-^ level in *bp-5 er-105* callus was lower compared with those in Col, *bp-5* and *er-105*(Fig. 2C). These results indicated that the function of H_2_O_2_ scavenge in *bp-5 er-105* double mutant may be defective during rapid growth stage of callus.

**Fig. 2.**
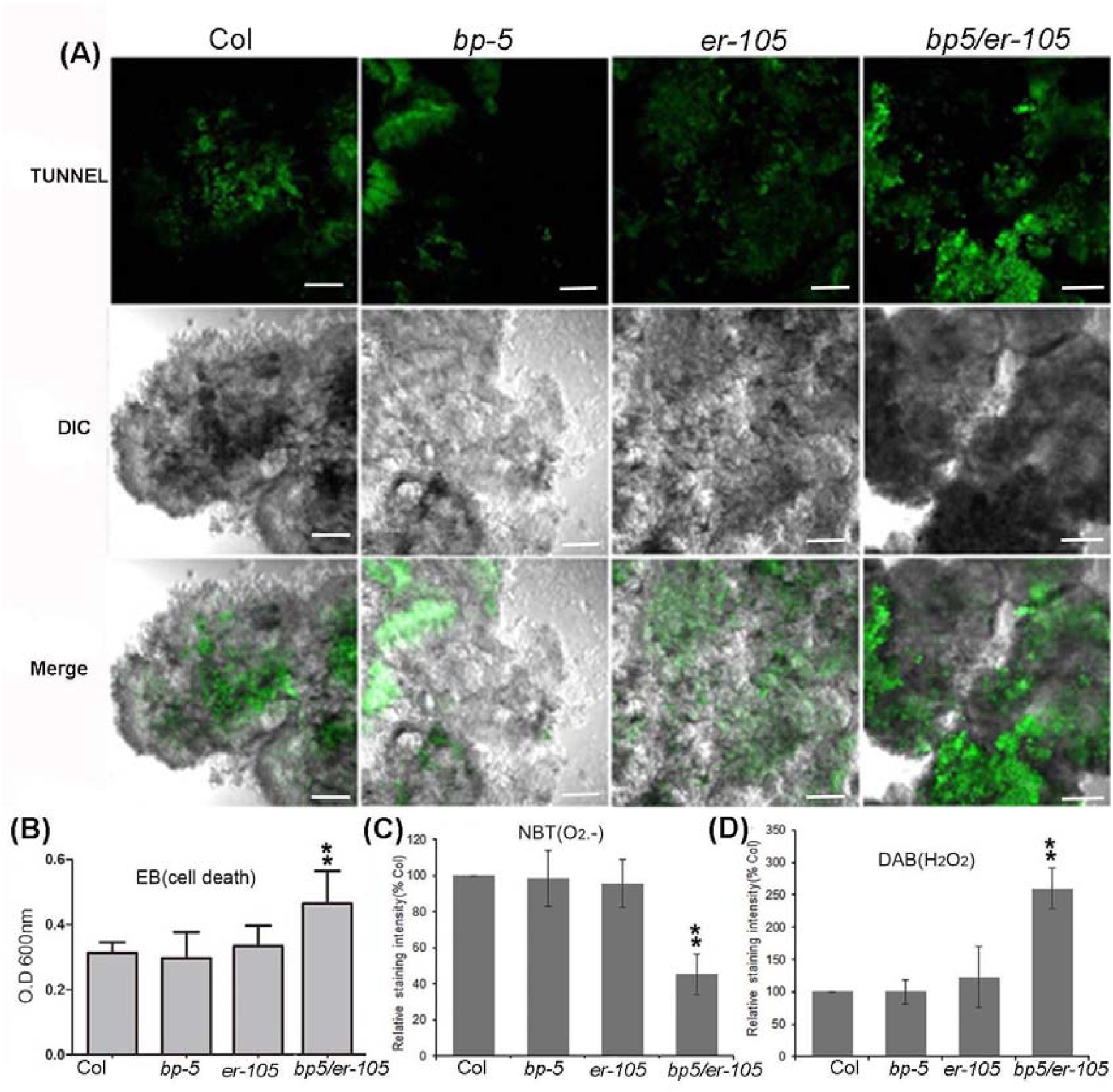
Comparison of cell death and endogenous O_2_^.-^ and H_2_O_2_ content in callus of *bp-5 er-105* with Col, *bp-5* and *er-105*. **(A)** Typical TUNEL assay fluorescence microscopy images of the 21-day calli and their phase-contrast microscopy (DIC) images. Scale bars, 200μm. **(B)** Evaluation of cell death in 21-day calli of Col, single mutants (*er-105* and *bp-5*) and double mutant (*bp-5 er-105*). The callus was stained with Evans blue (EB) as described in Methods. Cellular uptake of Evans blue was quantified by spectrophotometry at OD_600_ nm. Values represent means ± standard deviation (SD) (n=10; student’s t-test **p <0.001). **(C and D)** Quantification of nitroblue tetrazolium (NBT) and diaminobenzidine oxide (DAB staining intensity of Col, bp5, er105 and bp5 er105, respectively. The staining intensity of Col is given as 100%, of which the staining intensity of mutants was compared, respectively. Values represent means ± standard deviation (SD), (n=20, student’ s t-test **p <0.001).

### *BP* and *ER* promote expression of *AtPRX17* during callus growth

The class III perioxidases (E. C. 1.11.1.7, PRXs) in plant tissue play important roles in removal of excess H_2_O_2_ for maintaining a balance of ROS in tissue culture and the activities of peroxidase isoenzymes are important factors related to callus growth (Kay and Basile, 1987; Tournaire et al., 1996; Che et al., 2006).

We examine the patterns and activities of peroxidase isoenzymes in Col, *bp-5* and *er-105* and *bp-5 er-105* by a native polyacrylamide gel electrophoresis (PAGE) followed by in-gel 3, 3′-diaminobenzidine tetrahydrochloride (DAB) staining. Several bands corresponding to DAB-oxidizing active proteins were observed, among which one isoenzyme band was apparently weaker in *bp-5 er-105* callus than those in Col, and the corresponding single mutants *bp-5* and *er-105* callus (Fig. 3). These observations suggested that a specific peroxidase isoenzyme might be affected in *bp-5 er-105* during callus growth.

**Fig. 3.**
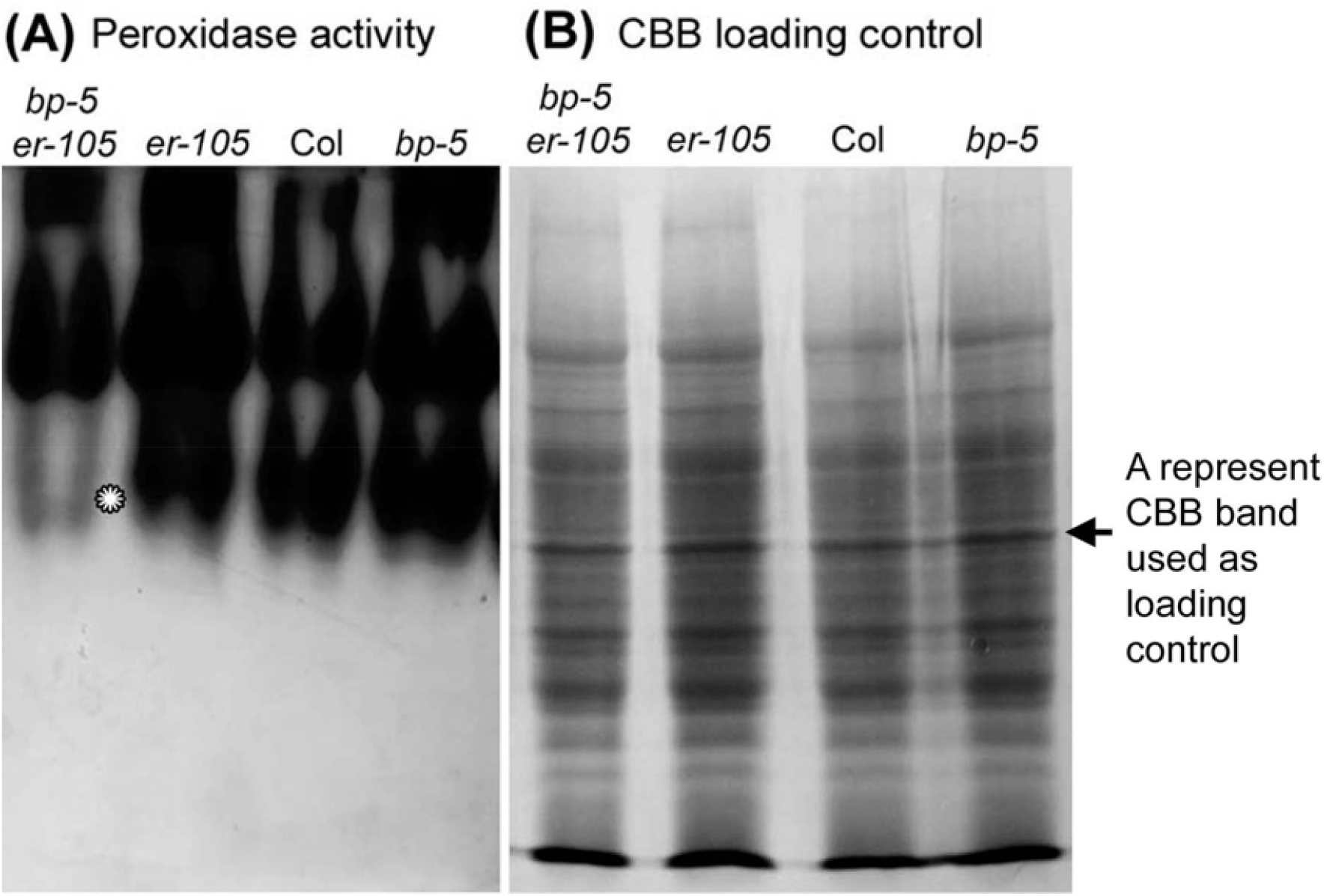
Isoenzyme analysis of peroxidases in developing callus of *bp-5 er-105, bp-5, er-105* and Col. **(A)** Native PAGE gels stained for peroxidase activity. Note that a band (asterisk indicating) in double mutant *bp-5 er-105* callus was weaker significantly in comparison with that of *bp-5, er-105* and Col callus, respectively. **(B)** Protein stained with Coomassie Brilliant Blue (CBB) was used as loading control.

To know if expression of some *PRX* genes in *bp-5 er-105* callus was altered at the transcription level, we re-examined expression levels of peroxidase genes in Arabidopsis callus according to microarray data published by previous studies (Che *et al*., 2002; Che *et al*., 2006). Among of the listed 73 genes of Arabidopsis class III PRXs, *peroxidase17* (At*PRX17*, At2g22420) and *peroxidase52* (At*PRX52*, At5g05340) were obviously up-regulated from 4 to 10 DAC during callus growth (Fig. S4), with the stage equivalent to that of rapid callus growth in our study (Fig. 1E). Reverse transcription polymerase chain reaction (RT-PCR) analysis revealed that expression of At*PRX52* in *bp-5 er-105* callus was not different from that in Col, *bp-5* and *er-105* callus (Fig. 4A), while expression of At*PRX17* was apparently down-regulated in the callus of *bp-5 er-105*, compared with that of Col, *bp-5* and *er-105* (Fig. 4A and B). At*PRX17* was selected for further study. A Salk T-DNA line (designated as *prx17*, SALK_034684c) was identified as homozygous by PCR analysis and there was no expression of At*PRX17* in the mutant callus (Fig S5). The growth rate of *prx17* callus was also apparently reduced in comparison with wild-type Col (Fig. 4E) and similar to that of *bp-5 er-105* callus (Fig. 1E). The patterns of PAGE showed that the dramatically reduced intensity of a band in *prx17* mutant callus was consistent with that in *bp-5 er-105* callus (Fig 4C). Thus, it is possible that the peroxidase isoenzyme At*PRX17* (isoPRX17) is defective in *bp-5 er-105* callus. To test the possibility of At*PRX17* expression regulated by *BP* and *ER*, we constructed a fusion with the At*PRX17* cDNA under the control of the *35S* promoter (*35S* _*pro*_*:: PRX17*), and introduced this fusion into the *bp-5 er-105* double mutant. A total of 12 independent transgenic plants were obtained, from which three homozygous lines for *35S* _*pro*_::At*PRX17* were subsequently identified for further characterization (Fig. 5A). The activities of peroxidases isoenzymes in cell cultures of these three transgenic lines were further examined, and the putative isoPRX17 band was rescued (Fig. 5B). Additionally, the callus growth and cell death defects in the 35S pro:PRX17/bp-5 er-105 transgenic plants were all rescued compared with those in Col and *bp-5 er-105* double mutant plants (Fig. 6A and B). These results indicated that *BP* and *ER* maintain normal callus growth possibly via promoting expression of At*PRX17*.

**Fig. 4.**
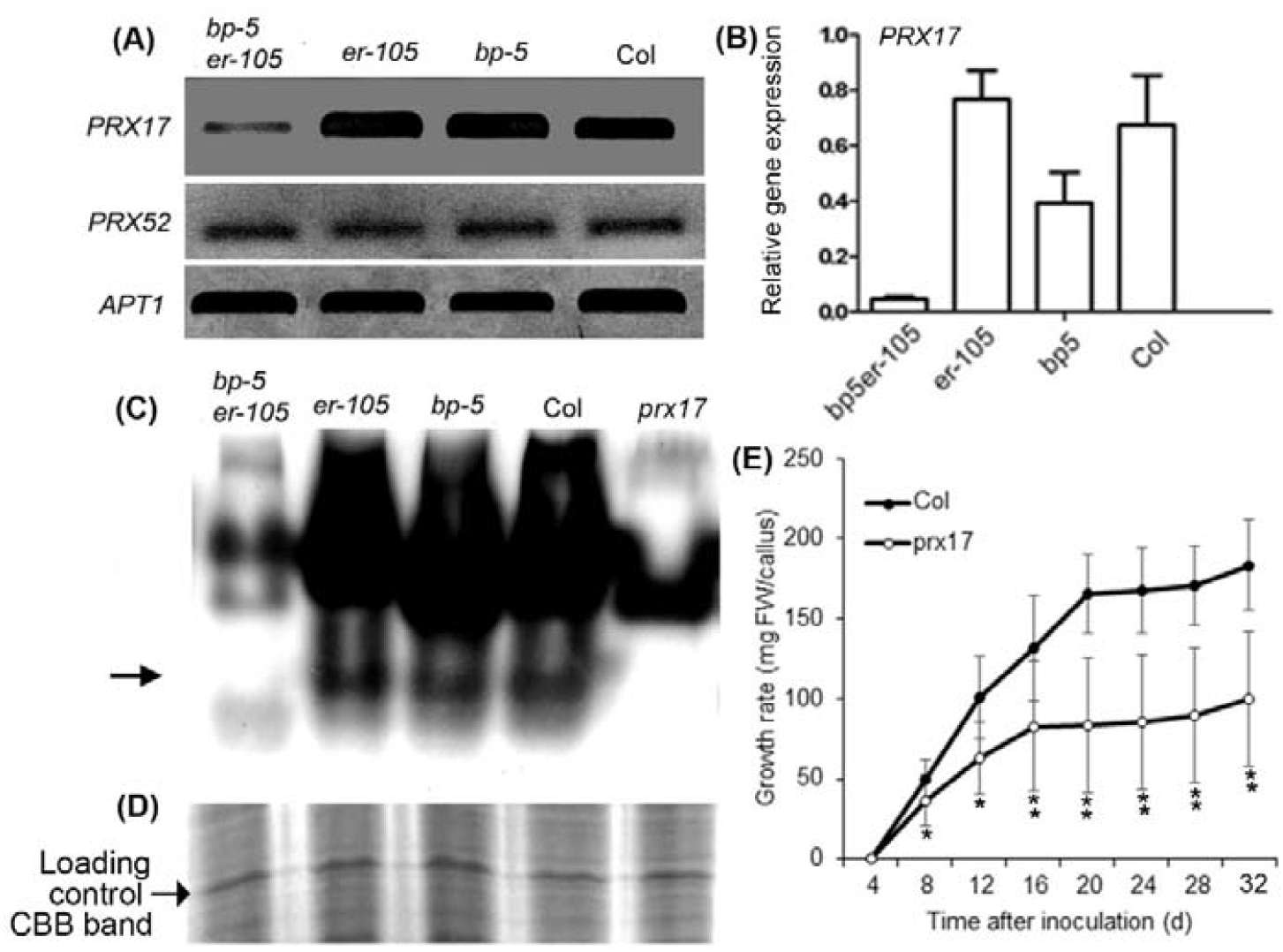
The absence of a peroxidase isoenzyme band of *bp-5 er-105* double mutant calli is consistent with that of *prx17* mutant calli. **(A)** The expression analysis of *PRX17* and *PRX52* genes in callus of different genotypes by RT-PCR. Note that expression of *PRX17* gene was apparently down-regulated in *bp-5 er-105* double mutant in comparison with that in *er-105, bp-5* and Col callus, respectively. Total RNA samples were extracted from callus grown on a new medium for14 days after subcultured. **(B)** qRT-PCR analysis of *PRX17* transcription levels in *bp-5er-105* in comparison with that in *er-105, bp-5* and Col calli, respectively. Values represent means±standard deviation (SD) (n=3) and results were consistent in at least three biological replicates. **(C)** Isoenzyme analysis of peroxidases in *bp-5 er-105, er-105, bp-5*, Col and *prx17* mutant callus by PAGE (50μg proteins/lane). Note that a band (arrows point) is absent in both *bp-5 er-105* and *prx17* in comparison with that in *er-105, bp-5* and Col, respectively. **(D)** A represent band in SDS-PAGE visualized by staining with Coomassie brilliant blue (CBB) was used as loading control. **(E)** Growth rate of *prx17* culture cells in comparison with that of Col. Error bars indicate the Standard deviation of the mean for three replications (n=16 calli for each time point; *p <0.05, **p <0.001, student’s t test).

**Fig. 5.**
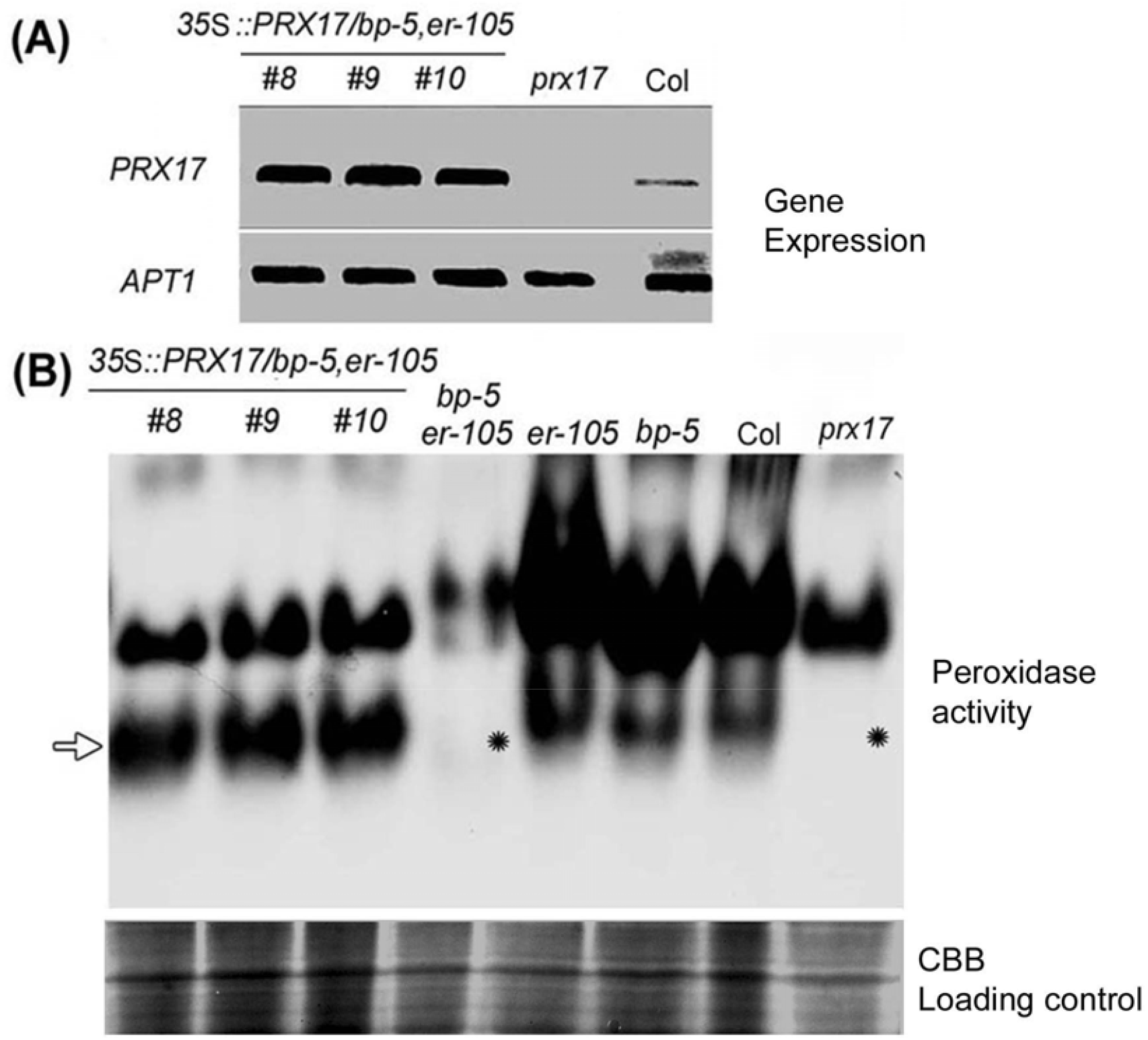
Substitution of *PRX*17 promoter with 35S promoter in *bp-5 er-105* double mutant could rescue the loss band of the peroxidase isoemzyme in the native PAEG gel. **(A)** RT-PCR analysis of *PRX*17 expression in *35S::PRX17/bp-5 er-105* transgenic lines(*bp-5 er-105* background). Total RNA samples were isolated from 14-day -old subculture callus. **(B)** Analysis of peroxidase isoenzymes in callus of *35S::PRX17/bp-5,er-105* transgenic line and the indicated mutants by PAGE (50μg proteins/lane). Noted that the lost band (asterisks indicate) of peroxidase isoemzymes in *bp-5 er-105* and *prx17* callus was rescued in the *35S::PRX17/bp-5 er-105* transgenic lines (e.g. line #8, #9 and #10) (arrow indicates). A represent band in SDS-PAGE visualized by staining with Coomassie brilliant blue (CBB) was used as loading control.

**Fig. 6.**
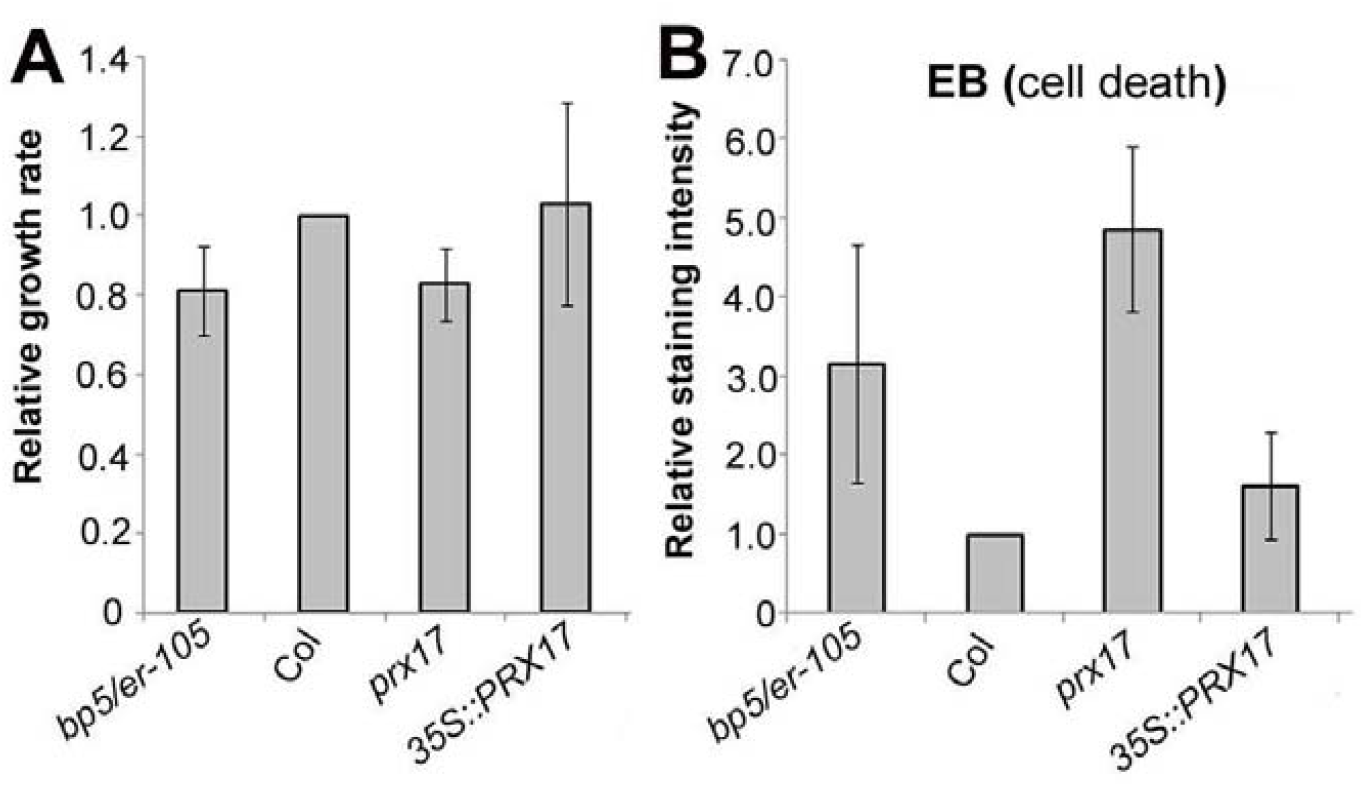
The growth rate of *prx17* mutant callus and *35S::PRX17* callus in comparison with that of *bp-5 er105* double mutant callus and wild-type Col callus, respectively. **(A)** Comparison of growth rate (expressed as an increase in callus fresh weight) of callus cultures derived from *bp5 er-105*, Col, *prx17, 35S:PRX17* root explants. 0.1g fresh weight callus of each indicating genotype was inoculated on the new CIM medium for 8 days and then measured the fresh weight. The increase in fresh weight of Col is given as 1.0, of which other mutants were compared, respectively. Values represent means ± standard deviation (SD) (n=15∼20). **(B)** Evaluation of cell death by Evans blue (EB) staining. Callus inoculated with new medium for 2 weeks was stained with Evans blue as described in Methods. Cellular uptake of Evans blue was quantified by spectrophotometry. Values represent means ± standard deviation (SD) (n=10 calli) of three experiments.

### BP binds to the TGAC motif in the At*PRX17* promoter

To investigate whether *BP* directly regulates At*PRX17* expression, we computationally searched for the nucleotide sequence similar to that of the published BP-binding *cis-*element from the genebank (www.arabidopsis.org) in At*PRX17* promoter (Smith et al., 2002). Three motifs consisting of TGACANCT (N =A or T) were identified in At*PRX17* promoter region between –1732 bp and –1359 bp from the translation initiation site, including TGACAACCT from –1712 to –1704, TGACAACT from –1680 to –1673, and TGACATCT from –1392 to –1385 (Fig. 7A). The interactions *in vitro* between the BP and the At*PRX 17* promoter region were further examined by electrophoretic mobility shift assays (EMSAs), using the recombinant BP proteins. The DNA fragment of the *AtPRX17* promoter region from –1732 to –1359 bp, which contains three BP-binding motifs, was used as probe. As shown in Figure 7A, a retarded band in the presence of the recombinant BP protein was observed. However, in the presence of excess amounts of the homologous unlabeled DNA fragment as competitor, the bund probe in the retarded band was apparently competed away. To further examine the function of BP in regulation of At*PRX17* expression *in vivo*, a transient expressional test was performed based on *agro*-infiltration of *Nicotiana benthamiana* leaves as described by the previous methods (Voinnet et al., 2003; Kane et al., 2007). In particular, tobacco leaves were co-infiltrated with two constructs: the reporter construct At*PRX17*_*pro*_*::GUS* and the effector construct *35S*_*pro*_*::BP* (Fig.7B). After infiltration and a 2-day recovery, the infiltrated leaves were collected for GUS staining, and GUS activity could reflect the At*PRX17* promoter activity. GUS staining revealed that reporter construct alone only resulted in relatively low At*PRX17* expression (Fig. 7C, left panel), whereas GUS staining apparently increased in tobacco leaves when co-infiltrated with both effector and reporter constructs (Fig. 7C, right panel). These results further supported that BP could directly regulate At*PRX17* gene expression. To provide more evidences that BP binds to the At*PRX17* promoter, we performed a chromatin imminoprecipitation (ChIP) assay using wild-type callus, and our results confirmed that BP could bind to At*PRX17* promoter *in vivo* (Fig. 7D).

**Fig. 7.**
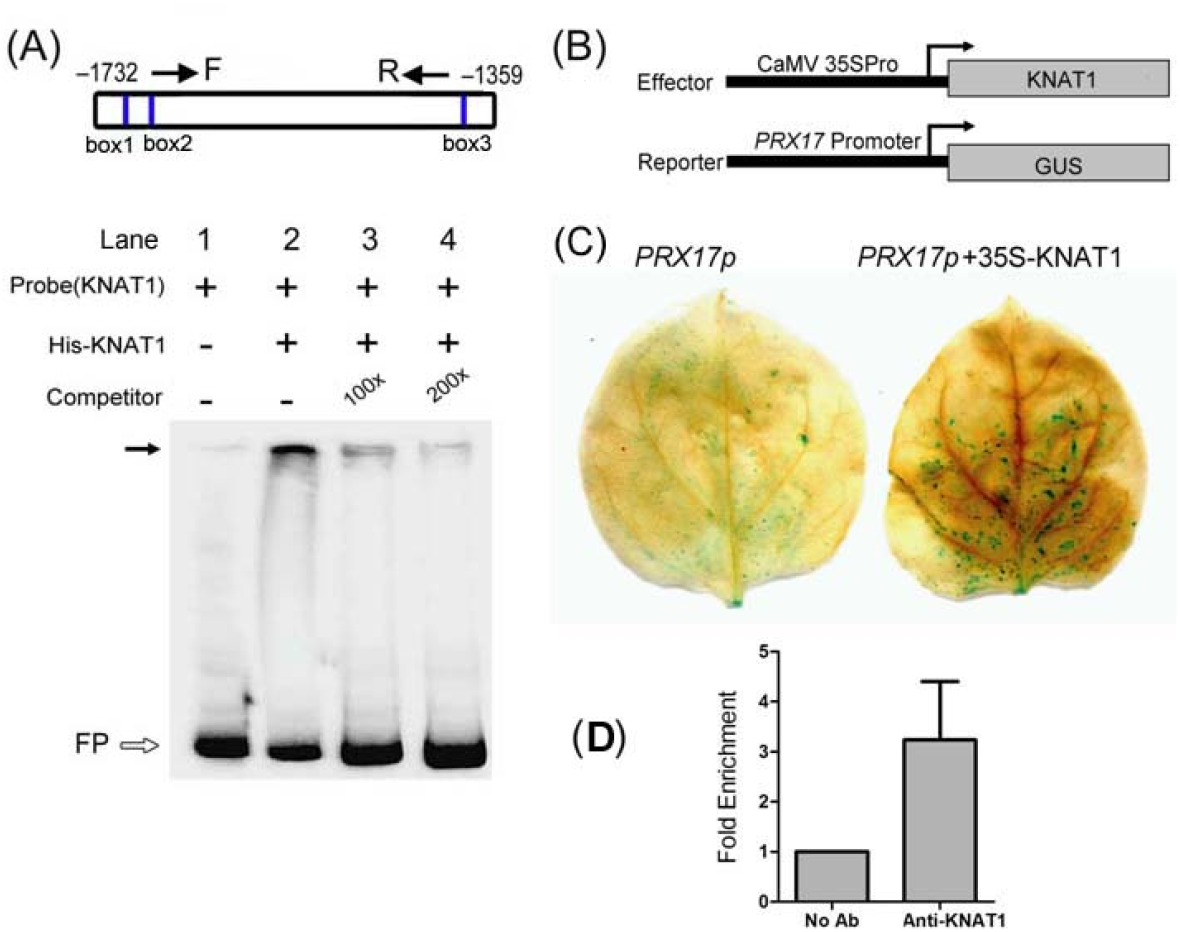
Interaction of KNAT1 protein with the promoter region of *PRX17*. **(A)** Electrophoretic mobility shift assay (EMSA) for KNAT1 binding to the *PRX17* gene promoter. The probe, biotin-labeled DNA corresponding to the *PRX17* gene promoter region (–1732 to –1359) from the site of initiation of translation and with three potential KNAT1 binding sites (box1, box2 and box3), was incubated in the absence (lane 1) and in the presence (lanes 2-4) of recombinant full-length KNAT1 proteins (His-KNAT1). As competitor DNA, homologous 100- and 200-fold excess of unlabeled DNA fragment was added to the reaction mixtures, respectively. The black arrows indicate shift band. The white arrows indicate free probe (FP). **(B)** Effector and Reporter constructs used in the transient assays. *pro35S*, Promoter from the *35S* gene of cauliflower mosaic virus; GUS, β-glucuronidase. **(C)** Effect of KNAT1 on *PRX17* promoter activity in vivo. *Nicotiana benthamiana* intact leaves were infiltrated with Agrobacterium strains carrying the reporter construct with or without effector constructs. pro*PRX17::GUS*, without effector; pro*PRX17::GUS*+pro*35S*:: *KNAT1 CDS*, with effector. Transactivation activity was detected by GUS staining assay. **(D)** Chromatin Immunoprecipitation (ChIP) assay of the Col callus showed that KNAT1 bound to *PRX17* promoter in *vivo* by qRT-PCR. Immunoprecipitation was performed with anti-KNAT1 antibody (Anti-KNAT1) or without antibody (no Ab). Primers (F+R) used in ChIP assay as shown in (A). Values represent means ± standard deviation (SD) (n= 3).

### The ER-downstream target *WRKY6* also directly regulates the At*PRX17* expression

ER protein is a membrane-bound leu-rich repeat receptor-like Ser/Thr kinase known as a pleiotropic regulator of multiple developmental and physiological processes and as a modulator to respond to environmental stimuli (Torii et al., 1996; Nanda et al., 2019). Previous studies show that ER regulates a set of *WRKY* transcription factor genes including *WRKY6, 15, 25, 33*, and *46* (Terpstra et al., 2010). To test whether ER modulates callus growth through WRKY genes, we first analyzed the expression of *WRKY* genes. Compared with Col, expression of *WRKY6* was apparently down-regulated in *er-105* and *bp-5 er-105* callus (Fig. 8A), whereas expression of several other WRKY genes was also analyzed, such as, *WRKY15, WRKY25, WRKY33* and *WRKY46*, but their expression was not significantly affected (Fig. 8A; Fig S6). Quantitative real-time PCR analysis demonstrated that expression of *WRKY6* in Col callus was gradually increased during 3 ∼9 DAC, and attained to the highest level at 9 DAC, coincident with expression pattern of *ER* during callus growth (Fig. 8B, compared to Fig. 1E). Thus, *WRKY6* was selected for further study.

**Fig. 8.**
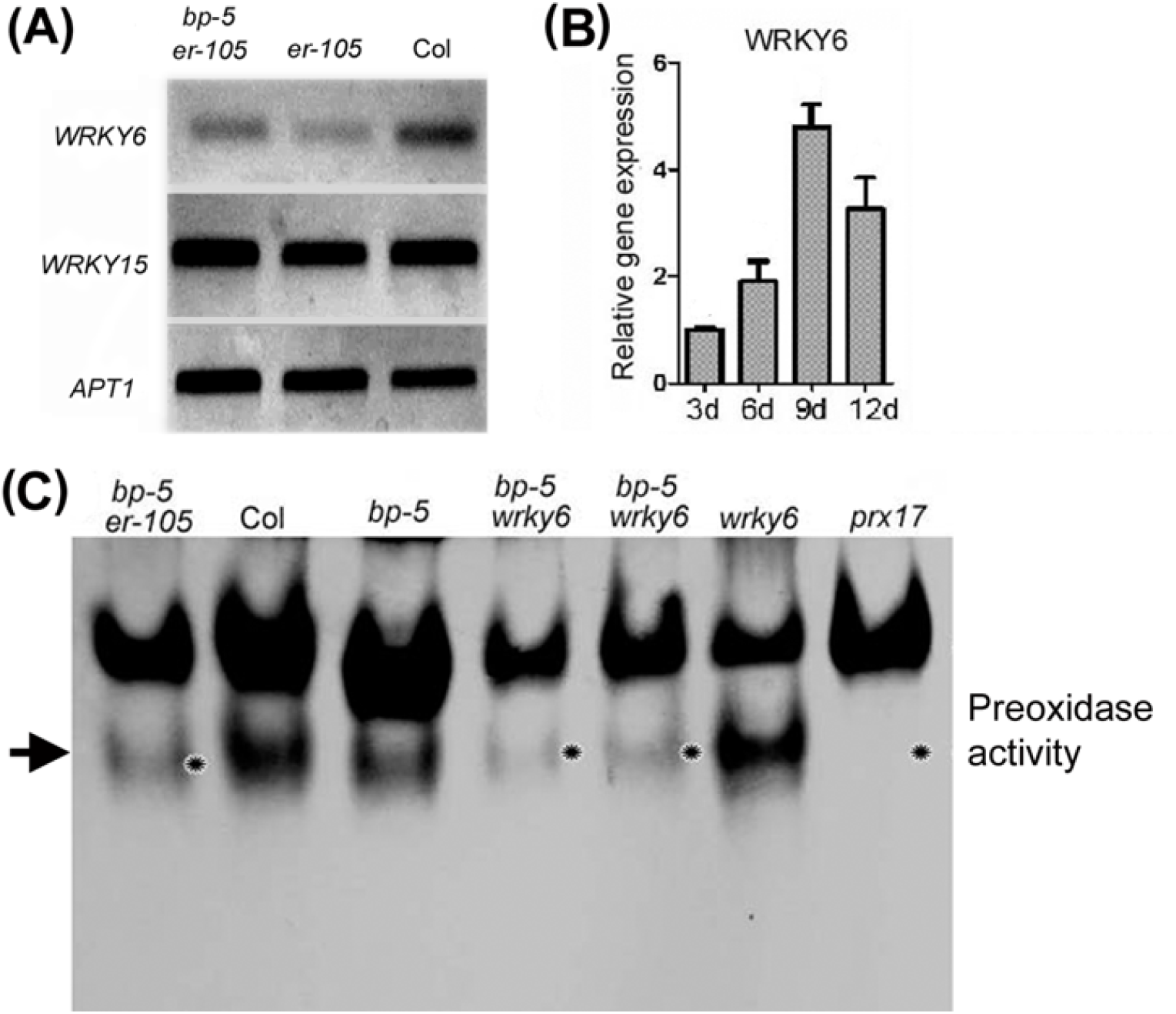
WRKY6 could together with BP and ER in regulating isoemzyme PRX17 activity in Arabidopsis callus. **(A)** RT-PCR analysis of *WRKY6* and *WRKY15* expression in the *er-105* and Col, respectively. *APT1* was used for the control. Total RNA samples from 14-day subculture old callus were isolated. **(B)** Relative transcript abundance changes of *WRKY6* gene in developing callus were detected using qRT-PCR. Callus were generated from root explants, and then inoculated on the new medium for 3, 6, 9 and 12 days. Values represent means ± standard deviation (SD) (n= 3). **(C)** Analysis of peroxidase isoenzymes in callus of indicated samples by PAGE (50μg proteins/lane). Noted that the lost band (asterisks indicate) of PRX17 isoenzyme in *bp-5 wrky6*, like that in *bp-5 er-105* callus (arrow indicates).

A mutant line with loss of *WRKY6* function, here referred to *wrky6*, was obtained (Fig. S7A and B), and used to construct *bp-5 wrky6* double mutant. Expression of *WRKY6* was severely reduced in *wrky6* and *bp-5 wrky6* double mutant (Fig S7C). The activity of the band corresponding to At*PRX17* in the *bp-5 wrky6* callus was apparently reduced, similar to that in *bp-5 er-105*, but markedly lower than those of the Col, *bp-5*, and *wrky6* (Fig. 8C). Furthermore, the growth rate of the *bp-5 wrky6* callus was significantly decreased compared with that of wild-type callus, but similar to that of the *bp-5 er-105* callus (Fig S8). These results indicated that *WRKY6* is involved in regulating the expression of At*PRX17*.

To further explore whether *WRKY6* also directly regulates theAt*PRX17*gene, we again examined the At*PRX17* promoter, trying to identify nucleotide sequence that WRKY6 binds to. Two previously reported WRKY-binding cis-motifs, TTGACC (Robatzek *et al*, 2002), were found in the At*PRX17* promoter between –1119 to –1114 and between –1005 and –1000 from the translation initiation site (Fig. 9A). The physical interaction between the WRKY6 and At*PRX17* promoter region was examined by EMSA as we have described above for the BP protein. The DNA fragment of the At*PRX17* promoter region from –1141 to –981 was used as probe. As shown in Figure 8A, a retarded band in the presence of the recombinant WRKY6 protein was observed. In the presence of excess amounts of the unlabeled competitor fragment, the amount of the labeled retarded complexes was obviously reduced (Fig. 9A). These results suggested that WRKY6 also specifically binds to the At*PRX17* promoter. To test the possible WRKY6 function in directly regulating At*PRX17*expression *in vivo*, we performed the transient assay through agro-infiltration of *Nicotiana benthamiana* leaves with the *35S*_*pro*_*::WRKY6* effector and *PRX17*_*pro*_*:: GUS* reporter constructs (Fig. 9B). GUS staining revealed that reporter construct alone only resulted in relatively low At*PRX17* expression (Fig. 9C, left panel), whereas GUS staining apparently increased in tobacco leaves when co-infiltrated with both effector and reporter constructs (Fig. 9C, right panel). Our results showed that similar to the *35S*_*pro*_*:: BP* effector, *35S*_*pro*_*::WRKY6* effector also enhanced the expression of At*PRX17*.

**Fig. 9.**
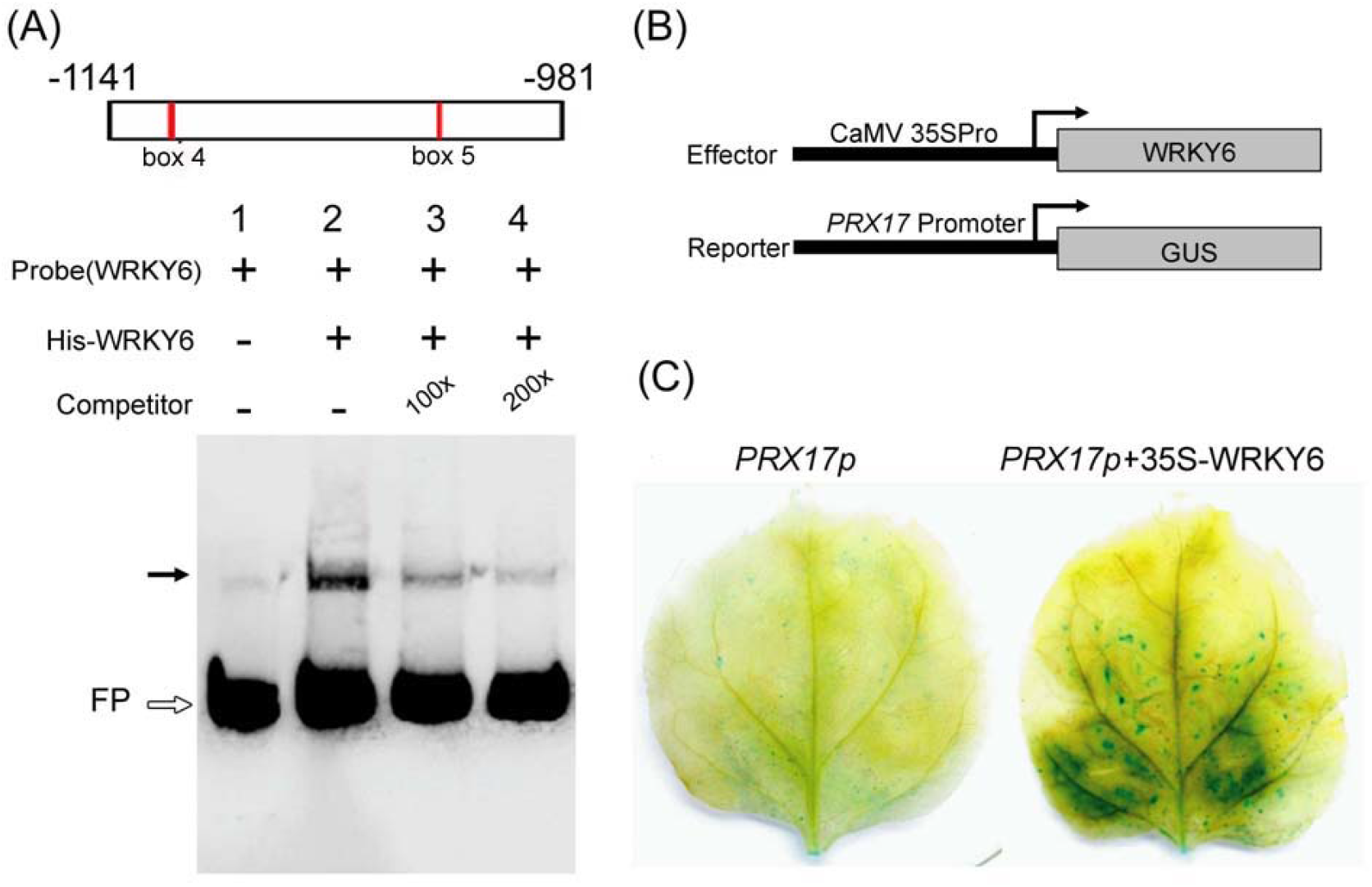
Interaction of WRKY6 protein with the promoter region of *PRX17*. **(A)** Electrophoretic mobility shift assay (EMSA) for WRKY6 binding to the *PRX17* gene promoter. The probe, biotin-labeled DNA corresponding to the *PRX17* gene promoter region (–1141 to –981) from the site of initiation of translation and two WRKY6 binding sites (box 4 and box 5), was incubated in the absence (lane 1) and in the presence (lanes 2-4) of recombinant full-length WRKY6 proteins. As competitor DNA, homologous 100- and 200-fold excess of unlabeled DNA fragment was added to the reaction mixtures, respectively. The black arrows indicate shift band. The white arrows indicate free probe (FP). **(B)** Effector and Reporter constructs used in the transient assays. *pro35S*, Promoter from the *35S* gene of cauliflower mosaic virus; GUS, β-glucuronidase. **(C)** Effect of WRKY6 on *PRX17* promoter activity in vivo. *Nicotiana benthamiana* intact leaves were infiltrated with Agrobacterium strains carrying the reporter construct with or without effector constructs. pro*PRX17::GUS*, without effector; pro*PRX17:: GUS*+pro*35S*::*WRKY6 CDS*, with effector. Transactivation activity was detected by GUS staining assay.

## Discussion

The class III peroxidases (PRXs) are a kind of plant-specific oxidoreductase that is involved in a broad range of physiological processes throughout the plant life cycle, including lignification, suberization, auxin catabolism, wound healing and defense against pathogen (Kay and Basile, 1987; Hiraga et al., 2001; Tognolli et al., 2002; Welinder et al., 2002; Passardi et al., 2006; Almagro et al., 2009; Cosio and Dunand, 2010; Herrero et al., 2013). Recent advances indicated that the processes directly or indirectly targeted by PRXs include gene expression, post-transcriptional reactions, and switching or tuning of metabolic pathways and other cell activities (Liebthal et al., 2018). However, little is known about the signal transduction for regulating expression of PRX genes. Previous studies show that the promoter region of peroxidase genes contains the transcription factor (TF) binding sequence. For example, AGL2 and /or WUS binding sites were detected in the promoter region of *PRX13, PRX30* and *PRX55* (Cosio and Dunand, 2010), but the specific function and precise role of these TFs remains unclear. In this study, we provide new evidences that BP together with WRKY6 directly regulate the expression of At*PRX17* in Arabidopsis callus by binding to its promoter region.

Callus is like the meristematic tissues, which is a self-renewing structure consisting of stem cells and their immediate daughters (Springer and Kohn, 1979; Xiao et al., 2020), but the mechanism on how to maintain this self-renewing structure is unclear. Reactive oxygen species (ROS) play an important role in maintaining plant cell proliferation (Vemoux et al., 2000; Wells et al., 2010). In Arabidopsis, two main ROS, superoxide (O_2_^.-^) and hydrogen peroxide exhibit distinct patterns of distribution in root tissues (Dunand et al., 2007). O_2_. ^-^ and H_2_O_2_ mainly accumulate in dividing and expanding cells in the meristem and elongation zones of root tips, respectively (Wells et al., 2010). Peroxidases have been suggested to be related with production (Mäder et al., 1980; Bolwell et al., 1995) and scavenging (Mehlhorn et al., 1996; Kvaratskhelia et al., 1997) of hydrogen peroxide and play an important role in the cell proliferation and senescence of higher plants (Abeles et al., 1988; Oh et al., 1997; Kay and Basile, 1987; Tsukagoshi et al., 2010).The class III peroxides multigene family in Arabidopsis genome show specific expression patterns in different developmental stages and organs (Tognolli et al., 2002). For example, in callus from Arabidopsis root explant, expression of *PRX73, PRX ATP21a, PRX27, PRXATP8a, PRX13a, PRXATP11a* and *ATP19a* were obviously down-regulated, while expression of *PRXATP12a* was up-regulated (Che et al., 2006). *PRX1*, which was mediated by *ABI3*, has been found to be specifically expressed in the embryo and aleurone layer during maturation and desiccation stage of development (Haslekås et al., 2003). Although these previous studies have found the appearance or disappearance of specific peroxidase isoforms during a particular process or in a particular localization (Loukii et al., 1999; Allison & Schultz, 2004), it is difficult to associate the band observed on an PAGE gel with a particular protein and define their roles in certain specific developmental processes, because protein purification of peroxidase isoenzymes is not straight forward as well as no obvious quantitative relationship exists between the transcript expression level and the protein activity (Dunand et al., 2003; Cosio and Dunand, 2009). In this study, either mutation of At*PRX17* gene in *prx17* mutant or impaired isoAtPRX17in double mutants *bp-5 er-105* or *bp-5 wrky6* resulted in an apparently decrease in the growth rate of callus (Fig 1E, Fig S8). Concentration of H_2_O_2_ in the isoPRX17 deficiency mutant callus was apparently higher than that in Col (Fig 1P, Fig 2C and D). Thus, we assumed that isoPRX17 could be essential in maintaining ROS homeostasis during Arabidopsis callus development. We further found that At*PRX17* could be specific in *BP*/*ER* pathway. *BP* has previously been shown to be redundant with *ER* to influence inflorescence stem and pedicel differentiation through local regulation of the level, or response to, a vasculature-associated growth inhibitory signal (Douglas et al., 2002; Douglas and Riggs, 2005). Recently *BP* and *ER* were found to involve in preventing precocious initiation of fiber differentiation during wood development (Vera-Sirera et al., 2019). However, which signaling pathway is involved in BP/ER pathway is still not clear. In this study, *BP* and *ER* are functionally redundant in promoting callus growth and inhibiting tissue browning. Analysis of the peroxidase isoenzyme patterns by a native PAGE gel indicated that double mutations in *BP* and *ER* genes resulted in deficiency of a PRX isoenzyme band, which is corresponding to AtPRX17isoenzyme (iso PRX17). Furthermore, the absence of isoPRX17 band in *bp-5 er-105* double mutant callus could be rescued in the *35S* _*pro*_*::PRX17/bp-5, er-105* transgenic lines. Thus, *BP* and *ER* could regulate expression of At*PRX17* gene by affecting its promoter activity. Although the BP recognition site is unknown, three binding sites for the KNOX gene BP/*KNAT1* (TGACAG(G/C)T)(Smith et al., 2002; Bolduc et al., 2012) are present at location –1732 bp to –1359 bp relative to the putative transcription start site in the promoter of At*PRX17*, suggesting that BP might directly bind to the At*PRX17* promoter region. Further EMSAs and transient assays using tobacco leaves also demonstrated that BP could directly bind to the At*PRX17* promoter region and affect its activity. However, the down-regulation of At*PRX17* transcript level was only observed in *bp-5 er-105* double mutant, but not in either *bp-5* or *er-105* single mutant. This implied that At*PRX17* could be a target of both BP and ER proteins. ER protein is a membrane-bound leu-rich repeat receptor-like Ser/Thr kinase (LRR-RLK; Torii et al., 1996; Van Zanten et al., 2009), but not a transcript factor (TF), it is impossible to regulate the expression of AtPRX17gene by directly binding to the promoter region. TFs regulated by ER have been reported (Terpstra et al., 2010). For example, *WRKY6* was suggested to act downstream of *ER* (Terpstra et al., 2010). In our study, two *WRKY6* binding sites (TTGACC) (Robatzek et al., 2002) close to one KNAT1 binding site (TGACATCT) were also observed in the promoter of *AtPRX17* from –1141bp to –981bp upstream of the putative transcription start site. In addition, WRKY6 protein bonded to the DNA segment of At*PRX17* promoter was also found with a much higher affinity *in vitro* and *vivo*. These observations suggest that the function of ER convergencely with BP to influence callus growth could be via WRKY6, which directly regulate expression of At*PRX17* gene.

Previous studies indicate that KNOX TFs in plant have degenerate binding sites and acquire specificity through cooperation with binding partners, as found in animals (Bolduc et al., 2009; Moens and Selleri, 2006). For example, KNOX proteins bind DNA as heterodimers with BELL proteins, another class of TALE HD protein (Bellaoui et al., 2001; Smith et al., 2002). KNOX and BELL share similar in vitro consensus binding sites, and their heterodimerization increases their affinity for DNA (Smith *et al*, 2002; Viola & Gonzalez, 2006). In this study, mutation of *BP* could cause inhibition of callus proliferation and At*PRX17* band deficiency, but this was only observed in an *er-105* or *wrky6* background. WRKY6 likely acts downstream of ER and as a cofactor of BP in regulating expression of At*PRX17* gene during callus development. Further studies should clarify the mechanism of how ER regulates *WRKY6* in Arabidopsis callus, and whether BP interacts with WRKY6 to form complex for regulating At*PRX17* in vivo.

## Supplementary data

**Fig. S1**. Callus was induced from Arabidopsis root explants.

**Fig. S2**. Callus induction from root explants of wild-type Col, single mutants *bp5* and *er-105* and double mutant *bp5*/*er-105* after 21 days on the callus induce medium.

**Fig. S3**. Growth of a double mutant *bp-1* and their single mutant L*er* (*BP/er*) and *bp/ER*, respectively, in comparison with their wild-type (Lan)..

**Fig. S4**. Time-resolved cluster analysis showing ratios of callus-developing related increases of Class III peroxidases in transcript abundances.

**Fig. S5**. Identification of homozygous insertion mutants at the PRX17 locus.

**Fig. S6**. RT-PCR analysis of WRKY25, WRKY33 and WRKY46 expression in the callus of *er-105* and Col, respectively.

**Fig. S7**. Identification of homozygous insertion mutants at the WRKY6 locus

**Fig. S8**. Comparison of growth rate of *bp-5 wrky6* double mutant callus with that of *bp-5 er-105* and *prx17* mutant callus, respectively.

**Table S1**. Oligonucleotide primers used in this study.

## Acknowledgments

The authors are indebted to Prof. Hai Huang for suggestions and Mr. Zhiping Zhang for EM works. This work was supported by the National Natural Science Foundation of China (31870850), the Strategic Pioneer Projects of CAS (XDB37020104), the China Manned Space Flight Technology project Chinese Space Station, the National natural fund joint fund project (U1738106).

## Author Contributions

JX carried out cell culture, data curation and analysis, BQ investigated gene expression; LW, YW and CM participated in investigation, HZ conceived of the study, participated in its design and coordination, and drafted the manuscript. All authors have read and agreed to the published version of the manuscript.

## References

Abeles FB, Dunn LJ, Morgens P, Callahan A, Dinterman RE, Schmidt J. 1988. Induction of 33-kD and 60-kD peroxidases during ethylene-induced senescence of cucumber cotyledons. Plant Physiology 87, 609–615.

Almagro L, LV Gómez Ros, Belchi-Navarro S, Bru R, Ros Barceló A, Pedreño MA.2009. ClassIII peroxidases in plant defence reactions. Journal of Experimental Botany 60, 377–390.

Allison, SD. and Schultz JC. 2004. Differential activity of peroxidase isozymes in response to wounding, gypsy moth, and plant hormones in northern red oak (Quercus rubra L.). Journal of Chemical Ecology 30,1363–1379.

Apel, K, Hirt H. 2004. Reactive oxygen species: Metabolism, oxidative stress, and signal transduction. Annual Review of Plant Biology 55, 373–399.

Baker CJ, Mock NM.1994. An improved method for monitoring cell death in cell suspension and disc assays using evans blue. Plant Cell, Tissue and Organ Culture 39, 7–12.

Bellaoui M, Pidkowich MS, Samach A, Kushalappa K, Kohalmi SE, Modrusan Z, Crosby WL, Haughn GW. 2001. The Arabidopsis BELL1 and KNOX TALE homeodomain proteins interact through a domain conserved between plants and animals. The Plant Cell 13, 2455–2470.

Basile, DV. 1980. A Possible Mode of Action for Morphoregulatory Hydroxyproline-Proteins. Bulletin of the Torrey Botanical Club 107, 325–338.

Bestwick, CS, Brown IR, Bennett, MHR, Mansfield JW. 1997. Localization of hydrogen peroxide accumulation during the hypersensitive reaction of lettuce cells to Pseudomonas syringae pv phaseolicola. The Plant Cell 9, 209–221.

Biswas, MS, Mano J. 2015. Lipid Peroxide-Derived Short-Chain Carbonyls Mediate Hydrogen Peroxide-Induced and Salt-Induced Programmed Cell Death in Plants. Plant Physiology 168, 885–898.

Bolduc N, Hake S. 2009. The maize trascription factor KNOTTED1 directly regulates the gibberellin catabolism gene ga2ox1.The Plant Cell 21, 1647–1658.

Bolduc N, Yilmaz A, Mejia-Guerra MK, Morohashi K, O’Connor D, Grotewold E, Hake S. 2012. Unraveling the KNOTTED1 regulatory network in maize meristems. Genes & Development 26,1685–1690.

Bolwell GP, Butt VS, Davies DR, Zimmerlin A. 1995. The origin of the oxidative burst in plants. Free Radical Research 23,517–532.

Breusegem FV, Dat JF. 2006. Reactive oxygen species in plant cell death. Plant Physiology 141,384–390.

Brewbaker JL, Upadhya MD, Mäkinen Y, MacDonald T. 1968. Isoenzyme polymorphism in flowering plants. III. Gel electrophoretic methods and applications. Physiologia Planturm 21,930–940.

Brown, JA, Li D, Alic M, Gold MH. 1993. Heat Shock Induction of Manganese Peroxidase Gene Transcription in Phanerochaete chrysosporium. Applied and Environmental Microbiology 59, 4295–4299.

Che P, Gingerich DJ, Lall S, Howell SH. 2002. Global and cytokinin-related gene expression changes during shoot development in Arabidopsis. The Plant Cell 14, 2771–2785.

Che P, Lall S, Nettleton D, Howell SH. 2006. Gene expression programs during shoot, root, and callus development in Arabidopsis tissue culture. Plant Physiology 141, 620–637.

Clough SJ, Bent AF. 1998. Floral dip: a simplified method 735–743.

Cosio, C and Dunand C. 2009. Specific functions of individual class III peroxidase genes. Journal of Experimental Botany 60, 391–408.

Cosio C, Dunand C. 2010. Transcriptome analysis of various flower and silique development stages indicates a set of class III peroxidase genes potentially involved in pod shattering in Arabidopsis thaliana. BMC Genomics 11, 528

Creissen CP, Edwards EA, Mullineaux PM. 1994. Glutathione reductase and ascorbate peroxidase. In Cause of photoxidative stress and amelioration of defense systems in plants, Foyer CH, Mullineaux PM (eds) pp 344–364. CRC Press, Boca Raton, USA

Daudi, A, Cheng Z, O’Brien JA, Mammarella N, Khan S, Ausubel FM, Bolwell GP. 2012. The apoplastic oxidative burst peroxidase in Arabidopsis is a major component of pattern-triggered immunity. The Plant Cell 4, 275–287.

Douglas SJ, Chuck G, Dengler RE, Pelecanda L, Riggs CD. 2002. BP and ERECTA regulate inflorescence architecture in Arabidopsis. The Plant Cell 14, 547–558

Douglas SJ, Riggs CD. 2005. Pedicel development in Arabidopsis thaliana: Contribution of vascular positioning and the role of the BREVIPEDICELLUS and ERECTA genes. Developmental Biology 284, 451–463.

Dunand, C, Meyer M De, Crevecoeur M, Penel C. 2003. Expression of a peroxidase gene in zucchini in relation with hypocotyl growth. Plant Physiology and Biochemistry 41, 805–811.

Dunand C, Crevecoeur M, Penel C. 2007. Distribution of superoxide and hydrogen peroxide in Arabidopsis root and their influence on root development:possible interaction with peroxidases. New Phytologist 174, 332–341.

Eulgem T and Somssich IE. 2007. Networks of WRKY transcription factors in defense signaling. Current Opinion in Plant Biology 10, 366–371.

Felipo-Benavent A, Urbez C, Blanco-Tourinan N, Serrano-Mislata A, Baumberger N, Achard P, Agusti J, Blazquez MA, Alabadi D. 2018. Regulation of xylem fiber differentiation by gibberellins through DELLA-KNAT1 interaction. Development 145, 1–7.

Goff, CW. 1975. Light and Electron-Microscopic Study of Peroxidase Localization in Onion Root Tip. American Journal of Botany 62, 280–291.

Habib D, Chaudhary MF, Zia M. 2014. The study of ascorbate peroxidase, catalase and peroxidase during in vitro regeneration of Argyrolobium roseum. Applied Biochemistry and Biotechnology 172, 1070–84.

Haslekås C,Viken MK, Grini PE, Nygaard V, Nordgard SH, Meza TJ, Aalen RB. 2003. Seed 1-cysteine peroxiredoxin antioxidants are not involved in dormancy, but contribute to inhibition of germination during stress. Plant Physiology 133,1148–1157.

Hay A, Tsiantis M. 2010. KNOX genes: versatile regulators of plant development and diversity. Development 137, 3153–3165.

Herrero J, Esteban-Carrasco A, Zapata JM. 2013. Looking for Arabidopsis thaliana peroxidases involved in lignin biosynthesis. Plant Physiology and Biochemistry 67,77–86.

Hord CLH, Suna YJ, Pillitteri LJ, Torii KU, Wang HC, Zhang SQ, Ma H. 2008. Regulation of Arabidopsis early anther development by the mitogen-activated protein kinases, MPK3 and MPK6, and the ERECTA and related receptor-like kinases. Molecular Plant 1: 645–658

Ikeuchi, M, Sugimoto, K IwaseA., 2013. Plant callus: mechanisms of induction and repression. The Plant cell 25, 3159–3173.

Kane NA, Agharbaoui Z, Diallo AO, Adam H, Tominaga Y, Ouellet F, Sarhan F. 2007. TaVRT2 represses transcription of the wheat vernalization gene TaVRN1. The Plant Journal 51, 670–680.

Kay LE, Basile DV. 1987. Specific peroxidase isoenzymes are correlated with organogenesis. Plant Physiology 84, 99–105.

Kvaratskhelia M,Winkel C, Thorneley RNF. 1997. Purification and characterization of a novel class III peroxidase isoenzyme from tea leaves. Plant Physiology 114:1237–1245

Laukkanen H, Haggman H, Kontunen-Soppela S, Hohtola A. 1999. Tissue browning of in vitro cultures of Scots pine: Role of peroxidase and polyphenol oxidase. Physiologia Planturm 106, 337–343.

Liebthal M, Maynard D, Dietz KJ. 2018. Peroxiredoxins and Redox Signaling in Plants. Antioxidants & Redox Signaling 28, 609–624.

Lincoln C, Long J, Yamaguchi J, Serikawa K, Hake S. 1994. A Knotted1-Like homeobox gene in Arabidopsis is expressed in the vegetative meristem and dramatically alters leaf morphology when overexpressed in transgenic plants. The Plant Cell 6, 1859–1876.

Loukili A, Limam F, Ayadi A, Boyer N, Ouelhazi L.1999. Purification and characterization of a neutral peroxidase induced by rubbing tomato internodes. Physiologia Planturm 105, 24–31.

Mäder M, Ungemach J, Schloss P. 1980. The role of peroxidase isoenzyme groups of Nicotiana tabacum in hydrogen peroxide formation. Planta 147, 467–470.

Mehlhorn H, Lelandais M, Korth HG, Foyer CH.1996. Ascorbate is the natural substrate for plant peroxidases. FEBS Letters 378, 203–206.

Mele G, Ori N, Sato Y, Hake S. 2003. The knotted1-like homeobox gene BREVIPEDICELLUS regulates cell differentiation by modulating metabolic pathways. Genes &Development 17, 2088–2093.

Milhinhos A, Vera-Sirera F, Blanco-Tourinan N, et al. 2019. SOBIR1/EVR prevents precocious initiation of fiber differentiation during wood development through a mechanism involving BP and ERECTA. Proceedings of the National Academy of Sciences of the United States of America 116, 18710–18716.

Moens CB, Selleri L. 2006. Hox cofactors in vertebrate development. Developmental Biology 291,193–206.

Murashige T, Skoog F. 1962. A revised medium for rapid growth and bioassays with tobacco tissue cultures. Physiologia Planturm 15, 473–497.

Nanda AK, El Habti A, Hocart CH, Masle J. 2019. ERECTA receptor-kinases play a key role in the appropriate timing of seed germination under changing salinity. Journal of Expermital Botany 70, 6417–6435.

Naton B, Ecke M, Hampp R.1992. Production of fertile hybirds by electrofusion of vacuolated and evacuolated tobacco mesophyll protoplasts. Plant Science 85, 197–208.

Oh SA, Park JH, Lee GI, Paek KH, Park SK, Nam HG.1997. Identification of three genetic loci controlling leaf senescence in Arabidopsis thaliana. The Plant Journal 12, 527–535.

Ori N, Eshed Y, Chuck G, Bowman JL, Hake S. 2000. Mechanisms that control knox gene expression in the Arabidopsis shoot. Development 127, 5523–5532.

Passardi F, Tognolli M, De Meyer M Penel, Dunand C. 2006. Two cell wall associated peroxidases from Arabidopsis influence root elongation. Planta 223, 965–974.

Qi B, Zheng HQ. 2013. Modulation of root-skewing responses by BP in Arabidopsis thaliana. The Plant Journal 76, 380–392.

Robatzek S, Somssich IE. 2002. Targets of AtWRKY6 regulation during plant senescence and pathogen defense. Genes & Development 16, 1139–1149.

Serikawa, KA, Martinez LA, Zambryski P. 1996. Three knotted1-like homeobox genes in Arabidopsis. Plant Molecular Biology 32, 673–683.

Sinha NR, Williams RE, Hake S. 1993. Overexpression of the maize homeobox gene, KNOTTED-1, causes a switch from determinated to indeterminate cell fates. Genes and Development 7, 787–795.

Shpak ED, Berthiaume CT, Hill EJ, Torii KU. 2003. Synergistic interaction of three ERECTA-family receptor-like kinases controls Arabidopsis organ growth and flower development by promoting cell proliferation. Development 131, 1491–1501.

Shen J, Xu G, Zheng HQ. 2015. Apoplastic barrier development and water transport in Zea mays seedling roots under salt and osmotic stresses. Protoplasma 252, 173–180.

Smith HM, Boschke I, Hake S. 2002. Selective interaction of plant homeodomain proteins mediates high DNA-binding affinity. Proceedings of the National Academy of Sciences of the United States of America 99, 9579–9584.

Springer WD, Green CE, Kohn KA. 1979. A histological examination of tissue culture initiation from immature embryos of Maize. Protoplasma 101, 269–281.

Tang, W, Newton RJ, Outhavong V. 2004. Exogenously added polyamines recover browning tissues into normal callus cultures and improve plant regeneration in pine. Physiologia Planturm 122, 386–395.

Terpstra IR, Snoek LB, Keurentjes JJB, Peeters AJM, van den Ackerveken G. 2010. Regulatory network identification by genetical genomics: signaling downstream of the Arabidopsis receptor-like kinase ERECTA. Plant Physiology 154, 1067–1078.

Tognolli M, Penel C, Greppin H Simon., 2002. Analysis and expression of the class III peroxidase large gene family in Arabidopsis thaliana. Gene 288,129–138.

Torii KU, Mitsukawa N, Oosumi T, Matsuura Y, Yokoyama R,Whittier RF, Komeda Y.1996. The Arabidopsis ERECTA gene encodes a putative receptor protein kinase with extracellular leucine-rich repeats.The Plant Cell 8, 735–746.

Tournaire, C, Kushnir S, Bauw G, Inze D, delaServe BT, Renaudin JP. 1996. A thiol protease and an anionic peroxidase are induced by lowering cytokinins during callus growth in Petunia. Plant Physiology 111, 159–168.

Tsukagoshi, H, Busch W, Benfey PN. 2010. Transcriptional regulation of ROS controls transition from proliferation to differentiation in the root. Cell 143, 606–616.

Venglat SP, Dumonceaux T, Rozwadowski K, Parnell L, Babic V, Keller W, Martienssen R, Selvaraj G, Datla R. 2002. The homeobox gene BREVIPEDICELLUS is a key regulator of inflorescence architecture in Arabidopsis. Proceedings of the National Academy of Sciences of the United States of America 99, 4730–4735.

Van Zanten M, Snoek LB, Proveniers MCG, Peeters AJM. 2009. The many functions of ERECTA. Trends in Plant Science 14, 214–218.

Vemoux T, Wilson RC, Seeley KA, Reichheld JP, Muroy S, Brown S, Maughan SC, Cobbett CS, van Montagu M, Inzé D et al. 2000. The ROOT MERISTEMLESS1/CADMIUM SENSITIVE2 gene defines a glutathione-dependent pathway involved in initiation and maintenance of cell division during postembryonic root development. The Plant Cell 12, 97–109.

Viola IL, Gonzalez DH. 2006. Interaction of the BELL-like proteinATH1 with DNA: Role of homeodomain residue 54 in specifying the different binding properties of BELL and KNOX proteins.Journal of Biological Chemistry 387, 31–40.

Voinnet O, Rivas S, Mestre P, Baulcombe D. 2003. An enhanced transient expression system in plants based on suppression of gene silencing by the p19 protein of tomato bushy stunt virus. The Plant Journal 33, 949–956.

Wakui K, Takahata Y, Kaizuma N. 1999. Scanning electron microscopy of desiccation-tolerant and sensitive microsporederived embryos of Brassica napus L. Plant Cell Reports 18, 595–600.

Wei N, Tan C, Qi B, Zhang Y, Xu GX, Zheng HQ. 2010. Changes in gravitational forces induce the modification of Arabidopsis thaliana silique pedicel positioning. Journal of Experimental Botany 61, 3875–3884.

Welinder KG, Justesen AF, Kjaersgard IVH, Jensen RB, Rasmussen SK, Jespersen HM, Duroux L. 2002. Structural diversity and transcription of class III peroxidases from Arabidopsis thaliana. European Journal of Biochemistry 269, 6063–6081.

Wells DM, Wilson MH, Bennett MJ. 2010. Feeling UPBEAT about growth: linking ROS gradients and cell proliferation. Developmental Cell 19, 644–646.

Woerlen N, Allam G, Popescu A, Corrigan L, Pautot V, Hepworth SR. 2017. Repression of BLADE-ON-PETIOLE genes by KNOX homeodomain protein BREVIPEDICELLUS is essential for differentiation of secondary xylem in Arabidopsis root. Planta 245,1079–1090.

Xiao Y, Li J, Zhang Y, Zhang X, Liu H, Qin Z, Chen B. 2020. Transcriptome analysis identifies genes involved in the somatic embryogenesis of Eucalyptus. BMC Genomics. 21, 803.

Zhang H, Chen J, Zhang F, Song Y. 2019. Transcriptome analysis of callus from melon. Gene 684,131–138.

Zhang K, Su J, Xu M, Zhou Z, Zhu X, Ma X, Hou J, Tan L, Zhu Z, Cai H et al. 2020 A common wild rice-derived BOC1 allele reduces callus browning in indica rice transformation. Nature Comnucation 11, 443.

Zhang Y, Wang L, Xie J, Zheng H. 2015. Differential protein expression profiling of Arabidopsis thaliana callus under microgravity on board the Chinese SZ-8 spacecraft. Planta 241, 475–488.

Zhao M, Li C, Ma X, Xia R, Chen J, Liu X, Ying P, Peng M, Wang J, Shi CL et al. 2020. KNOX protein KNAT1 regulates fruitlet abscission in litchi by repressing ethylene biosynthetic genes. Journal Experimental Botany 71, 4069–4082.

